# *Pf*Ago-based dual signal amplification biosensor for rapid and highly sensitive detection of alkaline phosphatase activity

**DOI:** 10.1101/2023.08.21.554052

**Authors:** Weikang Ke, Yuqing Qin, Bosheng Zhou, Yonggang Hu

**Affiliations:** National Key Laboratory of Agricultural Microbiology, College of Life Science and Technology, Huazhong Agricultural University, Wuhan 430070, P. R. China; Hubei Hongshan Laboratory, Wuhan 430070, P. R. China

**Author notes:** The authors wish it to be known that, in their opinion, the first two authors should be regarded as Joint First Authors.

**Keywords:** ALP activity, Biosensor, Human serum, *Pf*Ago

## Abstract

Developing rapid and highly sensitive methods for alkaline phosphatase (ALP) activity analysis is significant for the clinical diagnosis and treatment of diseases. Here, a *Pyrococcus furiosus* Argonaute (*Pf*Ago)-based biosensor is presented for ALP activity detection in which the ALP-catalyzed hydrolysis of 3’-phosphate-modified functional DNA activates the strand displacement amplification, and the amplicon -mediates the fluorescent reporter cleavage as a guide sequence of *Pf*Ago. Under the dual amplification mode of *Pf*Ago-catalyzed multiple-turnover cleavage activity and pre-amplification technology, the developed method was successfully applied in ALP activity analysis with a detection limit (LOD) of 0.0013 U L^−1^ (3σ) and a detection range of 0.0025 U L^−1^ to 1 U L^−1^ within 90 min. The *Pf*Ago-based method exhibits satisfactory analytic performance in the presence of the potential interferents and in complex human serum samples. The proposed method shows several advantages, such as rapid, highly sensitive, low-cost, and easy operation, and has great potential in disease evolution fundamental studies and clinical diagnosis applications.

## Introduction

Alkaline phosphatase (ALP) is widely found in living organisms and catalyzes the dephosphorylation reaction of biomolecules, such as nucleic acids and proteins(Millan 2006). For example, ALP participates in essential biological activities ranging from metabolism, molecule transduction, to cellular regulation(Niu et al. 2019). Consequently, ALP has been considered an important clinical biomarker for the diagnosis of diseases, such as cardiovascular diseases(Haarhaus et al. 2017), hepatobiliary diseases(Pratt and Kaplan 2000), leukemia(Neumann et al. 1974), osteoblastic bone tumor(Shimazaki et al. 2005), and diabetes(Tibi et al. 1988). Developing an accurate and sensitive ALP activity detection method is significant for uncovering the mechanism of ALP physiological functions underlying disease progression and meeting the demands for the clinical diagnosis and therapy of diseases.

Various methods have been developed for ALP activity detection, such as colorimetry(Niu et al. 2019), fluorimetry(Han et al. 2020, Niu et al. 2019), electrochemistry(Pan et al. 2020), and surface-enhanced Raman scattering(Sun et al.2020). Given the favorable analytical performance of the facile operation and the fast signal response and satisfying detection sensitivity of fluorescent methods, they have attracted remarkable attention for ALP activity analysis(Tang et al. 2019). For example, based on the ALP catalysis of the dephosphorylation reaction of 4-benzamio-2-(benzo[d]thiazol-2-yl)phenyl dihydrogen phosphate substrate to release fluorophore N-(3-(benzo[d]thiazol-2-yl)-4-hydroxyphenyl)benzamide, Yang et al. designed a “turn-on” ratiometric fluorescence method for ALP activity detection(Yangyang et al. 2020). Mao et. al. found that ALP can catalyze the hydrolysis of p-nitrophenol phosphate disodium salt to produce p-nitrophenol, which can quench the fluorescent signal of QDs by the inner filter effect. On the basis of this finding, they developed an indirect fluorescence method for ALP activity analysis.(Mao et al. 2019) According to ALP-catalyzed dephosphorylation reaction of 3’-phosphate termini ssDNA to generate 3’-hydroxyl ssDNA, Lee et al. presented a strand displacement amplification method, which is triggerred by the 3’-hydroxyl ssDNA products for ALP activity analysis. The ALP activity is proportional to the fluorescence signal obtained by the interaction between the amplicon and SYBR green I(Lee, Park and Park 2018). Lai et al. proposed a DNAzyme-regulated CRISPR/12a-assisted method for ALP activity detection(Lai et al. 2022). They used a Cu^2+^-dependent DNAzyme to catalyze the cleavage of a long DNA sequence, which can bind with Cas12a-crRNA to activate the trans-cleavage activity of CRISPR/Cas12a toward the fluorescence probe, leading to the loss of the trans-cleavage reaction of CRISPR/Cas12a. Meanwhile, the DNAzyme activity is inhibited because of the free Cu^2+^’s in the presence of pyrophosphate via the formation of Cu^2+^-pyrophosphate complexation. In this case, Cu^2+^ could be released from the Cu^2+^-pyrophosphate complexation after adding ALP through pyrophosphate hydrolysis, and the Cu^2+^ activates the DNAzyme-CRISPR/12a cascade reaction. Thus, the fluorescence intensity is related to ALP activity. Recently, Wang et al. presented a CRISPR/Cas13a-based dual-amplification method for ALP activity analysis(Wang et al. 2021b). In their work, a 5’-phosphate-modified dsDNA was catalyzed to generate 5’-hydroxyl dsDNA in the presence of ALP. The 5’-hydroxyl dsDNA triggers the T7 polymerase mediated transcription amplification, and the amplicon activates the trans-cleavage activity of CRISPR/Cas13a toward the fluorescence probe to release the fluorescent signal. The ALP activity could be detected by the change in fluorescent signal. The dual-amplification mode could realize the highly specific and sensitive analysis of ALP activity with a LOD of 0.006 U L^−1^. However, the CRISPR/Cas system requires a guide RNA and the recognition of a PAM sequence, which results in its low adaptability and high cost, limiting its applications for the on-site monitoring of ALP activity(Lin et al. 2022). Developing a simple, rapid, and highly sensitive platform for ALP activity analysis is still preferred.

Argonaute proteins (Agos) exist in nearly all organisms and play a crucial role in RNA or DNA interference pathways to defend the host against invasive nucleic acid molecules(Meister 2013, Swarts et al. 2014). The Ago system has been described as an emerging programmable nucleic acid biotechnology, which can employ a short ssDNA as the guide sequence mediated by the complementary ssDNA cleavage(Qin, Li and Hu 2022). Agos-based biosensors have been successfully used for nucleic acid target detection, including SARS-CoV-2(Wang et al. 2021a, Xun et al. 2021), miRNAs(Lin et al. 2022, Shin et al. 2020), pathogenic bacteria(Li et al. 2023), and human cancer(Liu et al. 2021, Song et al. 2020). In this work, we developed a *Pyrococcus furiosus* Argonaute (*Pf*Ago)-mediated biosensor for the rapid and sensitive detection of ALP activity. The proposed method shows several advantages, such as rapidity, high sensitivity, eco-friendlinrdd, and low cost, and has a promising potential for application inr clinical diagnosis and disease therapy.

## Experimental Section

### Materials

ALP and ATP were purchased from BBI Life Sciences Co., Ltd. (Shanghai, China). DNA sequence includes S1 (5’-AGG ACC TCT AAC CTC AGC AAC ATT ACT GCA GAT TCA CAA CAT TTT TTT AAT GTT GTG-P-3’), T1-FQ (5’-FAM-CGC ACC AGG ACC TCT AAC CTC AGG TGC G-BHQ-3’), T2-FQ (5’-FAM-CAT ACC TCG AGG ACC TC-BHQ-3’), and gDNA (5’-TGA GGT TAG AGG TCC T-3’) were synthesized by Sangon Biotech Co., Ltd. (Shanghai, China). Klenow Fragment, Nb.BbvCI, and NEB Buffer 2 were obtained from New England BioLabs Inc. (Beijing, China). HEPES, NaCl, and other chemical reagents used in this work were obtained from Sinopharm Chemical Reagent Co., Ltd. (Shanghai, China). The reagents dissolution and reaction system preparation were used by ultrapure H_2_O (18.20 MΩ) with an ultrapure H_2_O system from Pall Co., Ltd (Washington, USA).

### Preparation of *Pf*Ago protein

Expression and purification of *Pf*Ago as follows(Swarts et al. 2015): 1) The synthetic gene sequence of *Pf*Ago was cloned into pET28a with a His tag in the N-terminal, named as *Pf*Ago-pET28a. 2) The recombined plasmid was transformed into *E*.*coli* BL21 (DE3) and the transformant was cultivated with the LB medium (containing 50 μg mL^−1^ kanamycin) in a shaker incubator (37°C, 180 rpm). When the cell concentration is about OD_600_=0.60–0.80, IPTG (0.1 mmol L^−1^) was added into the medium and incubated at 16°C for 20 h. 3) The cells were obtained by centrifuged (7 000 g, 5 min), and using lysis buffer (20 mmol L^−1^ Tris-HCl, 500 mmol L^−1^ NaCl, 2 mmol L^−1^ MnCl_2_, pH 8.0) to resuspend the cells. The cells were broken by a pressure-crushing system (1 200 bar, 20 min). Then, the crushing liquid was removed by centrifuging (12 000 g, 45 min). The Ni-NTA sefinose resin kit was used for *Pf*Ago protein purification with elution buffer (20 mmol L^−1^ Tris-HCl, 500 mmol L^−1^ NaCl, 2 mmol L^−1^ MnCl_2_, and different imidazole concentrations, pH 8.0). The obtained protein was analyzed by SDS-PAGE and Western Blot. The purified *Pf*Ago was stored at −20°C for usage.

### Detection of ALP activity

The effect of 3’P-S1, T1-FQ, *Pf*Ago, and *Pf*Ago-catalyzed reaction time for ALP detection was investigated, as follows. 1) ALP catalyzed dephosphorylation system (total 10 mL reaction system): 3’P-S1 (0.1, 0.2, 0.3, 0.4, 0.5, 0.6 μmol L^−1^), AP buffer (1×), and ALP (1 U L^−1^) was incubated at 37°C for 15 min. Then, the system was heated at 75°C for 2 min to deactivate the ALP activity. 2) Strand Displacement Amplification system (total 15 mL reaction system): dNTP (0.83 mmol L^−1^), Klenow Fragment (50 U mL^−1^), Nb.BbvCI (200 U mL^−1^), and NEB Buffer 2 (1×) were incubated at 37°C for 30 min. 3) *Pf*Ago-mediated cleavage process (total 50 mL reaction system): *Pf*Ago (0.12, 0.16, 0.20, 0.24, 0.28 mg mL^−1^), T1-FQ (0.14, 0.29, 0.43, 0.57, 0.71 μmol L^−1^), and *Pf*Ago buffer (8 mmol L^−1^ HEPES, 80 mmol L^−1^ NaCl, 0.80 mmol L^−1^ MnCl_2_, pH 8.0) were incubated at 95°C for different time (20, 25, 35, 40, 45, and 50 min). The fluorescent signal was recorded using Fluoroskan Ascent FL Multifunctional Microplate Reader (Thermo Fisher Scientific, Waltham, MA, USA) with excitation wavelength at 485 nm and emission wavelength at 520 nm. Under the optimum conditions, the detection performance of the method was evaluated by adding different ALP concentrations (0.001, 0.0025, 0.005, 0.01, 0.05, 0.1, 0.5, and 1 U L^−1^) into the system.

### The performance investigation of the *Pf*Ago-based method

The inhibition model was investigated by adding Na_3_VO_4_ (0, 25, 50, 100, 150, 200, 250, 300 μmol L^−1^) and ATP (0, 5, 10, 25, 50, 75, 100 μmol L^−1^) into the ALP activity analysis system, respectively. The reaction process is described above. The effect of potential interferents included BSA (100 mg mL^−1^), Cys (3 mmol L^−1^), ALP (1 U L^−1^), the heated ALP (1 U L^−1^, 75°C, 5 min), glucose oxidase (10 U L^−1^), and HRP (10 U L^−1^) on the ALP analysis method were studied. The reaction conditions as described above.

To investigate the potential applications in clinical diagnosis, the *Pf*Ago-based biosensor was used to evaluate the ALP level of human serum samples. Firstly, the samples were diluted 500 times and filtered using the DNA Mini columns with a nucleic acid-affinity membrane (Omega Bio-tek.) to decrease or remove the nucleic acid in samples. The ALP activity of serum samples was analyzed by the proposed biosensor and the obtained result was compared with the testing from the hospital.

### The multi-turnover analysis of *Pf*Ago

Firstly, *Pf*Ago (0.05 μmol L^−1^) was incubated with gDNA (0.05 μmol L^−1^) in *Pf*Ago buffer (8 mmol L^−1^ HEPES, 80 mmol L^−1^ NaCl, 0.8 mmol L^−1^ MnCl_2_, pH 8) at 55°C for 30 min for the formation of gDNA-*Pf*Ago complex. Then, different concentrations of ssDNA-FQ (0.05, 0.25, and 0.5 μmol L^−1^) were added into the gDNA-*Pf*Ago mixture incubated at 95°C for different times (0, 5, 10, 15, 20, 25, and 30 min), respectively. The fluorescent intensity readout procedure was followed as above shown.

**Scheme 1.**
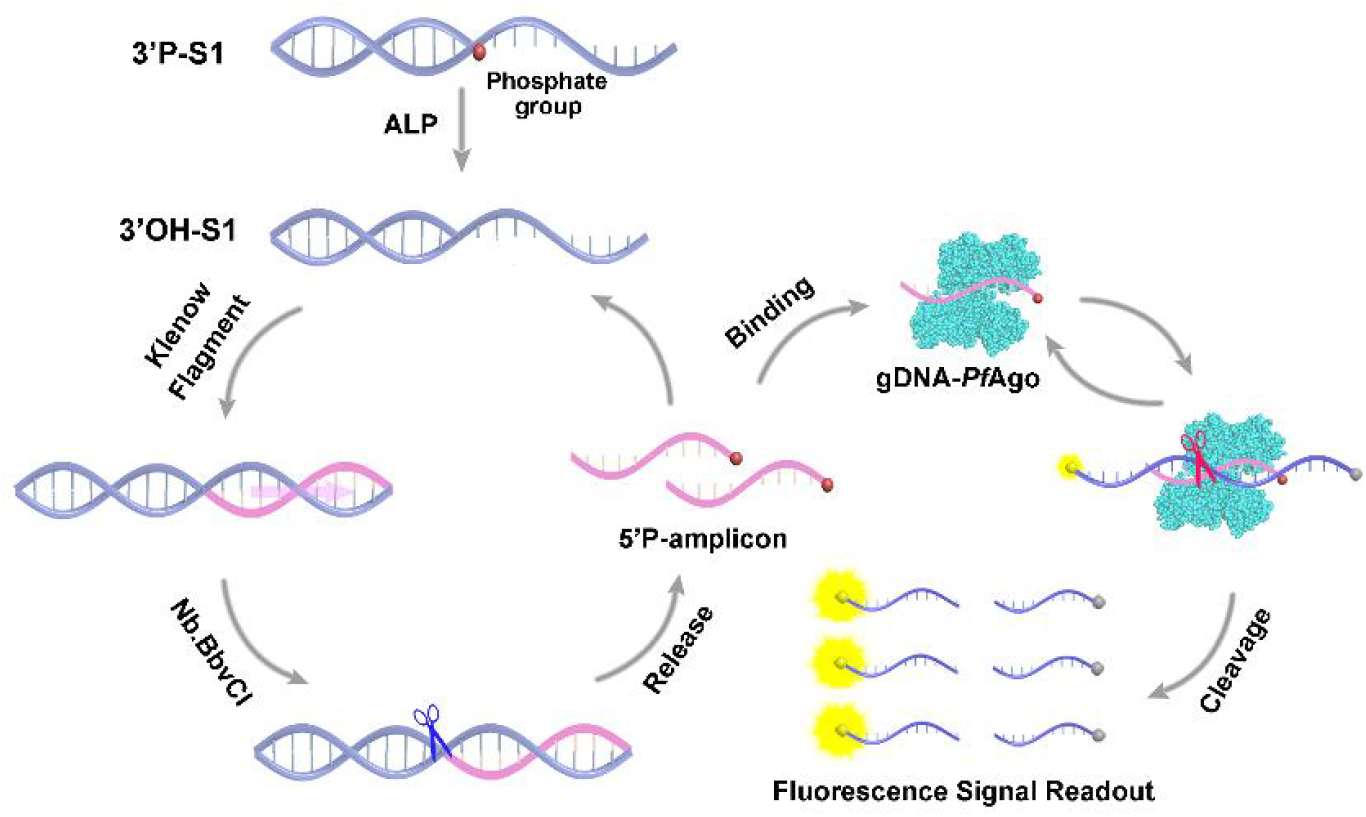
*Pf*Ago-based biosensor used for ALP activity detection.

## Results and Discussion

As shown in Figure S1, a band of approximately 90 kDa was observed in the SDS-PAGE, indicating that *Pf*Ago was successfully obtained(Swarts et al. 2015). *Pf*Ago employs 5’-phosphate termini ssDNA (5’P-ssDNA) as the guide sequence (gDNA) to form the gDNA-*Pf*Ago complex to activate the cleavage of the complementary ssDNA target through base pairing (Figure S2). According to their programmable nucleic acid property, a *Pf*Ago-based strategy was developed for ALP activity analysis. As shown in Scheme 1, an incomplete dsDNA was designed with the 3’-phosphate termini named 3’P-S1. In the presence of ALP, the phosphate group of 3’P-S1 was hydrolyzed to produce 3’-hydroxyl termini S1 (3’OH-S1), which could be recognized by Klenow Fragment polymerase and induced by the Klenow Fragment/Nb.BbvCI-mediated strand displacement amplification (SDA). The obtained amplicon with the 5’-phosphate termini acted as gDNA to activate the *Pf*Ago-catalyzed cleavage of the ssDNA probe (T1-FQ) labeled with a fluorophore–quencher pair, and the fluorescence signal was recorded by a multifunctional microplate reader. As shown in Figure 1A, PAGE analysis showed that a band appeared, which corresponds to 3’P-S1 (Lane 1). No obvious band change was observed with the addition of the SDA reagent into the above system (Lane 3) because the Klenow Fragment lacks the 5’-3’ exonuclease activity, and its polymerase I activity cannot recognize the 3’-phosphate group and further amplify the DNA sequence(Henner, Grunberg and Haseltine 1983). In the presence of ALP, the phosphate group of 3’ P -S1 was removed, and the obtained 3’-OH S1 was recognized by the Klenow Fragment and activated the SDA reaction. As expected, an obvious amplicon band appeared at the lower site analyzed by PAGE (Lane 4). A weak band appeared only in the presence of the ssDNA-FQ fluorescence reporter (T1-FQ) due to the quenched reaction of the fluorophore–quencher pair labeled (Lane 2). The band of the amplicon disappeared, and a band appeared at the molecular weight between S1 and ssDNA-FQ when the reporter was added into the ALP-mediated SDA system (Lane 5) because T1-FQ and the amplicon formed a dsDNA through base-pairing. An obvious cleavage band appeared and the dsDNA and S1 bands faded With the addition of *Pf*Ago (Lane 6), illustrating that *Pf*Ago could employ the amplicon as the guide to catalyze the T1-FQ cleavage. These results demonstrate that the proposed *Pf*Ago-based biosensor could be used for ALP detection. To verify the strategy for the *Pf*Ago-mediated dual-signal amplification, commercial 5’-OH ssDNA, commercial 5’-P ssDNA, or amplicon was incubated with *Pf*Ago/F1-FQ, and the fluorescence signal from the F1-FQ cleavage was recorded by a fluorescence microplate reader. As shown in Figure 1B, without ALP, the obtained signal intensity was similar to the negative sample (a). Using the commercial 5’-P ssDNA (d) as the guide of *Pf*Ago instead of 5’-OH ssDNA (c) generated a detectable signal. An obvious fluorescent signal was observed when ALP-induced amplicon was added into the *Pf*Ago/T1-FQ system (e). These results demonstrate that the amplicon obtained from SDA could activate the *Pf*Ago-catalyzed T1-FQ cleavage reaction to realize the highly sensitive detection of ALP activity.

**Figure 1.**
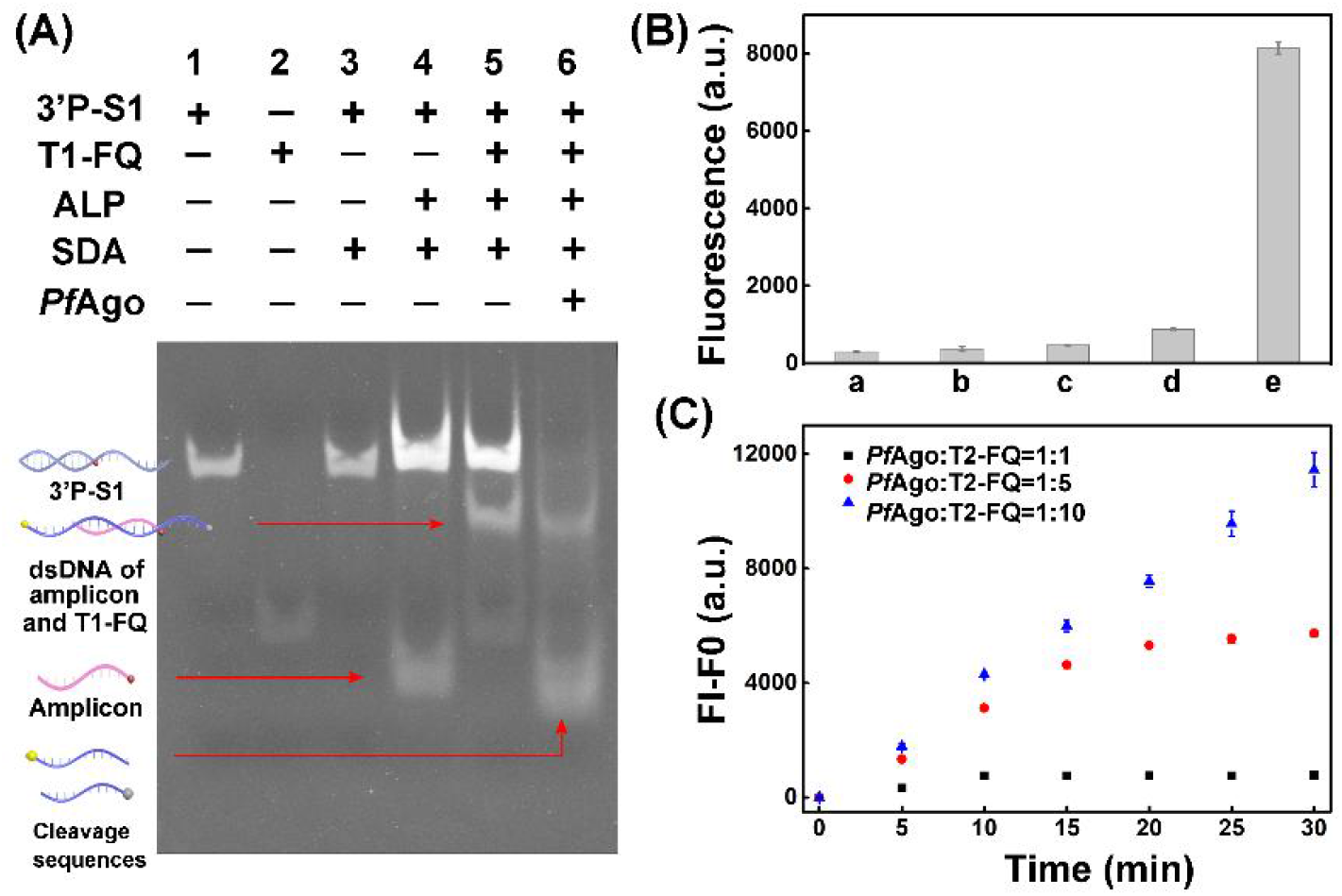
(A) PAGE analysis of the *Pf*Ago-based biosensor for ALP activity analysis. Lane 1: 3’P-S1; Lane 2: T1-FQ; Lane 3: 3’P-S1 + Klenow Fragment + Nb.BbvCI; Lane 4: ALP + 3’P-S1 + Klenow Fragment + Nb.BbvCI; Lane 5: ALP + 3’P-S1 + Klenow Fragment + Nb.BbvCI + T1-FQ; Lane 6: ALP + 3’P-S1 + Klenow Fragment + Nb.BbvCI + *Pf*Ago + T1-FQ. (B) Fluorescence intensity results of different detection systems. a: T1-FQ; b: 3’P-S1 + Klenow Fragment + Nb.BbvCI + *Pf*Ago + T1-FQ; c: Commercial 5’OH-gDNA + *Pf*Ago + T1-FQ; d: Commercial 5’P-gDNA + *Pf*Ago + T1-FQ; e: ALP + 3’P-S1 + Klenow Fragment + Nb.BbvCI + T1-FQ; Lane 6: ALP + 3’P-S1 + Klenow Fragment + Nb.BbvCI + PfAgo + T1-FQ. (C) The cleavage property of *Pf*Ago was studied under the conditions of both single turnover (0.050 μmol L^−1^ T2-FQ), and multi-turnover (0.25 μmol L^−1^, and 0.50 μmol L^−1^ T2-FQ).

The turnover catalysis functions of *Pf*Ago were studied by incubating *Pf*Ago, gDNA, and T2-FQ in various molecular ratios at 75 °C for 0–30 min (Figure 1C). At the 1:1:1 molecular ratio of *Pf*Ago, gDNA, and T2-FQ (0.050:0.050:0.050 μmol L^−1^), the fluorescent intensity gradually increased and reached equilibrium within 10 min. The fluorescent signal was enhanced by 6.92-fold within 20 min at the molar ratio of 1:1:5 (0.050:0.050:0.25 μmol L^−1^) compared with that in the initial system. Compared with that at the 1:1:1 molecular ration of *Pf*Ago, gDNA, and T2-FQ, a 14.77-fold fluorescent intensity increase in the *Pf*Ago-catalyzed T2-FQ cleavage was observed in 30 min with the addition of T2-FQ into the reaction at the concentration ratio of 1:1:10 (0.050:0.050:0.50 μmol L^−1^). And the signal continuously increased at the initial velocity. These results demonstrate the multiple-turnover activity of *Pf*Ago for the nucleic acid target cleavage, which is consistent with the findings of a previously reported work(Swarts et al. 2015).

The effect of S1 (0.10–0.60 μmol L^−1^), T1-FQ (0.14–0.71 μmol L^−1^), *Pf*Ago (0.12–0.28 mg mL^−1^), and *Pf*Ago catalysis reaction time (20–50 min) on ALP activity detection was investigated (Figure 2). Under optimum conditions (Dephosphorylation system: 1 U L^−1^ ALP, 1×AP buffer, and 0.4 μmol L^−1^ S1, incubation at 37 °C for 15 min, and heating at 75 °C for 2 min; The SDA system: 50 U mL^−1^ Klenow Fragment, 200 U mL^−1^ Nb.BbvCI, 0.83 mmol L^−1^ dNTP, and 1×NEB buffer 2, incubation at 37 °C for 30 min; The *Pf*Ago cleavage system: 0.2 mg mL^−1^ *Pf*Ago and 0.43 μmol L^−1^ T1-FQ, incubation at 95 °C for 45 min), the fluorescent intensity gradually increased with the addition of ALP. A good linear relationship existed between the fluorescence intensity and ALP activity in the range of 0.0025–1 U L^−1^, fitting the following equation (Figure 3A): *F* = 4247*C* + 138 (R^2^ = 0.99),

**Figure 2.**
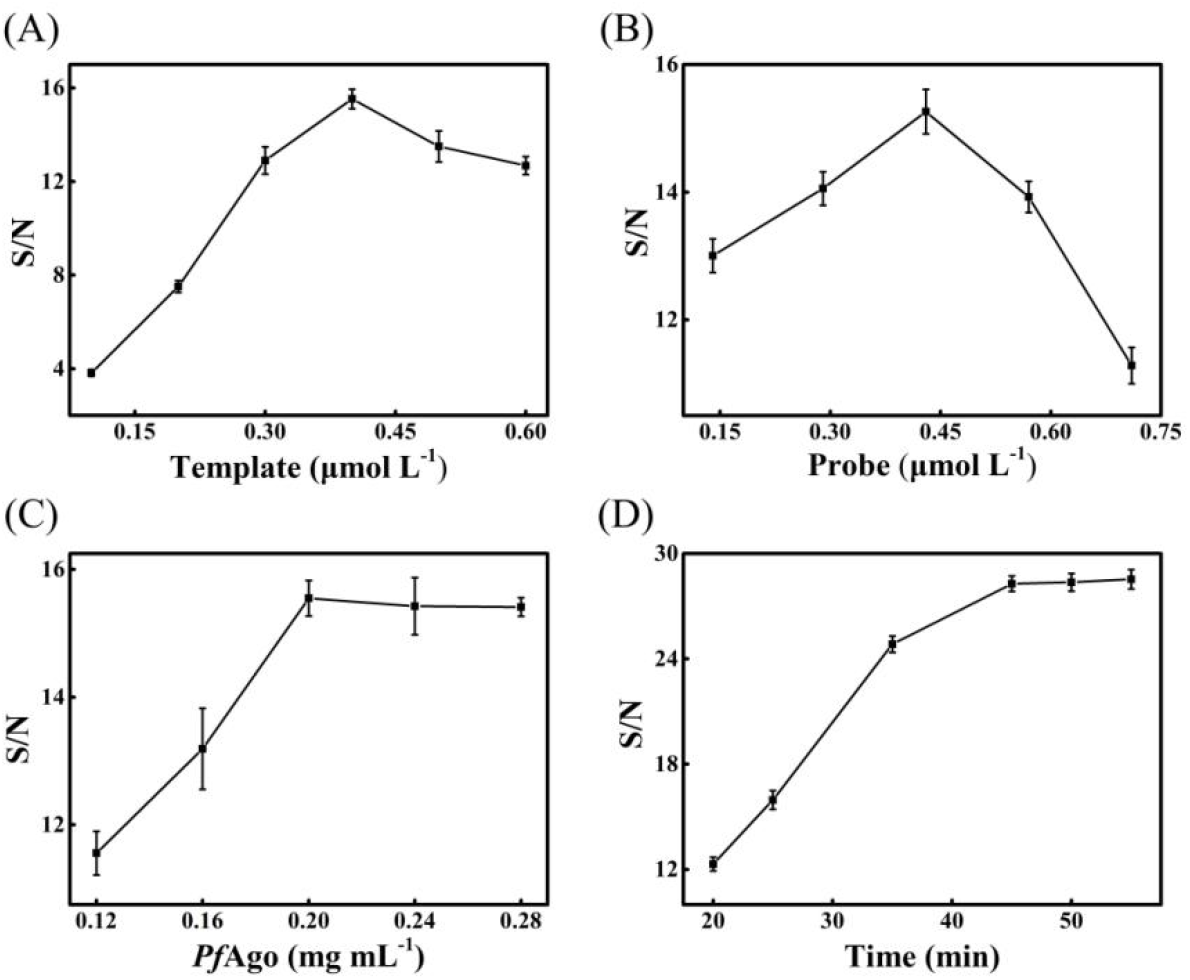
The effect of Template (A), Probe (B), *Pf*Ago (C), Time (D) for the *Pf*Ago-based sensing system for ALP detection. Conditions: The dephosphorylation system: 1×AP buffer, 1 U L^−1^ ALP and 0.10–0.60 μmol L^−1^ 3’P-S1 (A) were incubated at 37°C for 15 min and heated at 75°C for 2 min; The SDA system: 0.83 mmol L^−1^ dNTP, 50 U mL^−1^ Klenow Fragment, 200 U mL^−1^ Nb.BbvCI and 1×NEB buffer 2 were incubated at 37°C for 30 min; The cleavage system: 0.14–0.71 μmol L^−1^ F1-FQ (B) and 0.12–0.28 mg mL^−1^ *Pf*Ago (C) were incubated at 95°C for different 20-50 min (D). The error bars represent the SE of the means. The data represent means ± SE (n = 3).

**Figure 3.**
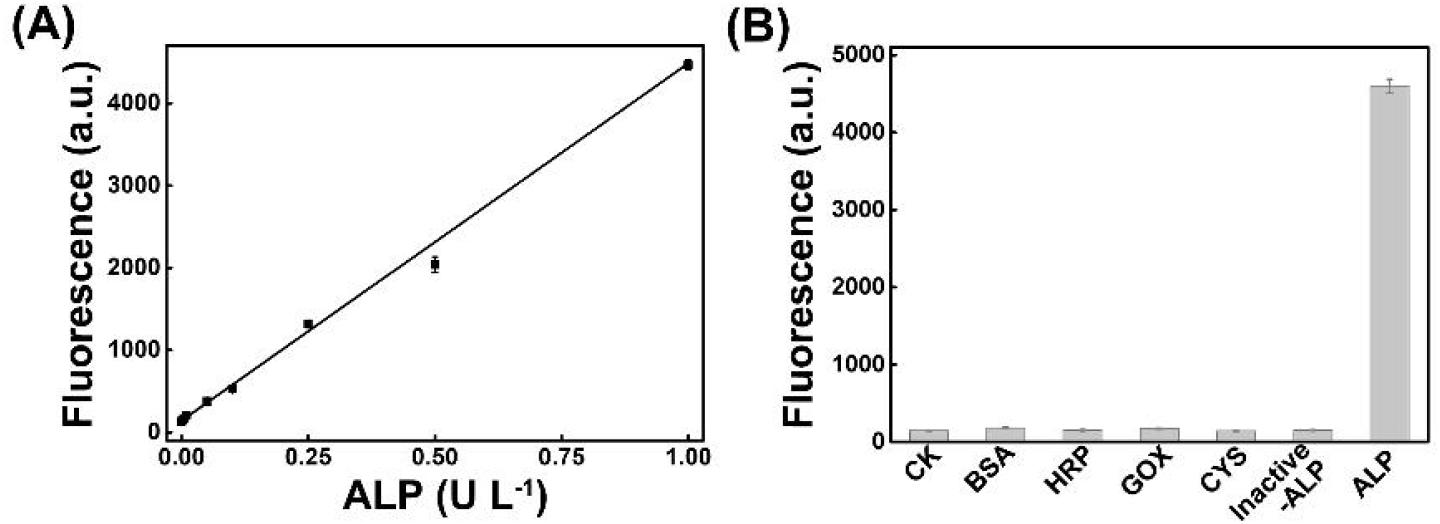
(A) *Pf*Ago-based strategy used for ALP activity detection. (B) The effect of the potential interferents on the sensing system. Conditions: The dephosphorylation system: 1×AP buffer and 0.4 μmol L^−1^ 3’P-S1 were incubated at 37°C for 15 min and heated at 75°C for 2 min; The SDA system: 0.83 mmol L^−1^ dNTP, 50 U mL^−1^ Klenow Fragment, 200 U mL^−1^ Nb.BbvCI and 1×NEB buffer 2 were incubated at 37°C for 30 min; The cleavage system: 0.2 mg mL^−1^ *Pf*Ago and 0.43 μmol L^−1^ F1-FQ were incubated at 95°C for 45 min. ALP activity includes 0.0025, 0.0050, 0.010, 0.050, 0.10, 0.25, 0.50 and 1 U L^−1^ (A); 1 U L^−1^ ALP, 100 mg mL^−1^ BSA, 10 U L^−1^ HRP, 10 U L^−1^ glucose oxidase (GOX), 3 mmol L^−1^ Cysteine, and 1 U L^−1^ inactive ALP (B). The error bars represent the SE of the means. The data represent means ± SE (n = 3).

**Figure 4.**
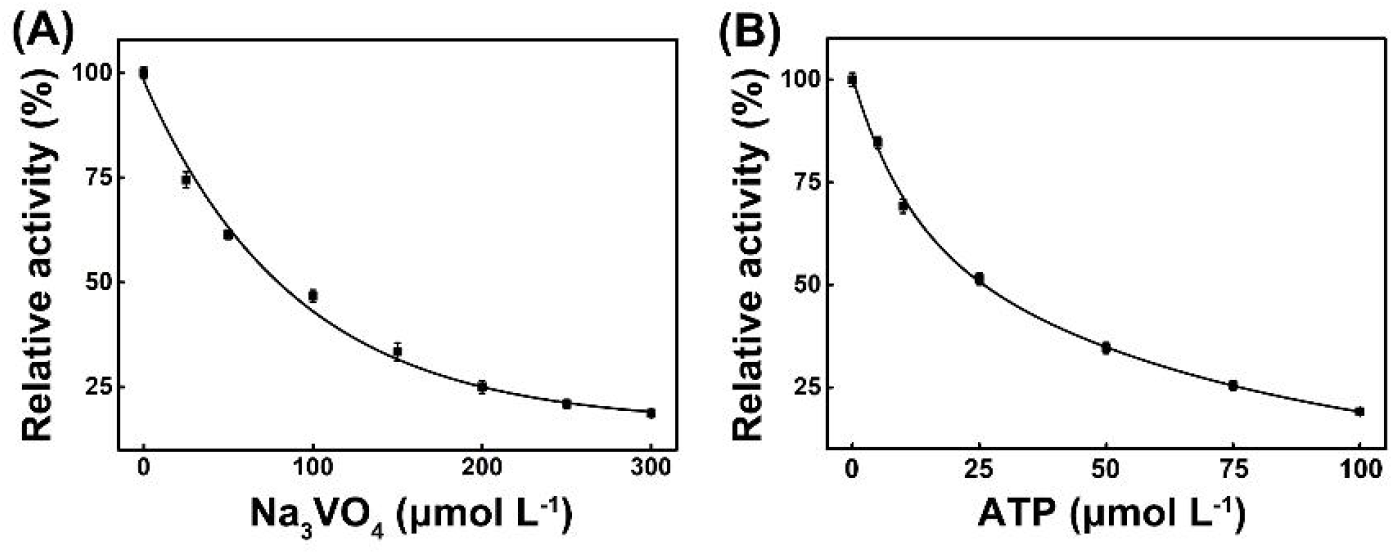
Inhibition of ALP activity by Na_3_VO_4_ (A) and ATP (B). Conditions: The dephosphorylation system: 1 U L^−1^ ALP, 1×AP buffer, 0.40 μmol L^−1^ 3’P-S1, and different concentration inhibitors were incubated at 37°C for 15 min and heated at 75°C for 2 min; The SDA system: 0.83 mmol L^−1^ dNTP, 50 U mL^−1^ Klenow Fragment, 200 U mL^−1^ Nb.BbvCI, and 1×NEB buffer 2 were incubated at 37°C for 30 min; The cleavage system: 0.2 mg mL^−1^ *Pf*Ago and 0.43 μmol L^−1^ T1-FQ were incubated at 95°C for 45 min. The error bars represent the SE of the means. The data is presented as means ± SE (n = 3).

where *F* is the relative fluorescence intensity, and *C* is the ALP activity (U L^−1^). The LOD was calculated at 0.0013 U L^−1^ (3σ/k), which is comparable to that in reported works (Table 1). The developed strategy realized the rapid and highly sensitive detection of ALP activity, which can be attributed to signal amplification steps of SDA-based pre-amplification and the multiple turnover activity of the *Pf*Ago-catalyzed T1-FQ cleavage. The influence of the potential interferents, such as BSA, glucose oxidase, HRP, cysteine, and heated-inactive ALP, on the *Pf*Ago-based biosensor for ALP activity analysis was investigated. As shown in Figure 3B, a large fluorescence intensity was recorded in the presence of ALP, and the interferents could not induce the detectable signal. These results show the good anti-interference capacity of the proposed biosensor for ALP activity analysis.

**Table 1.**
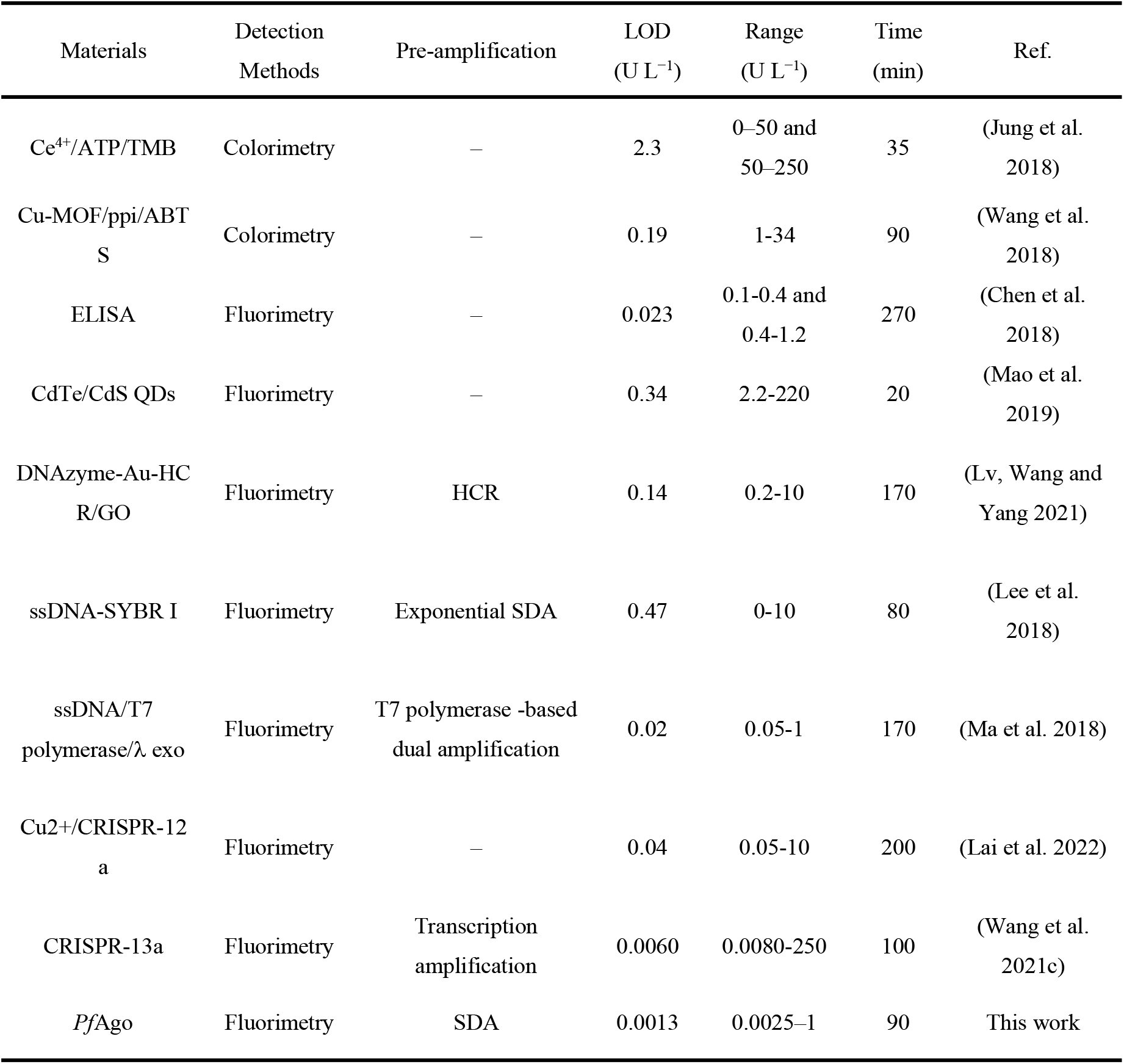
Comparison of the proposed method and the methods in the reported studies

Given that the ALP level is associated with several human diseases, screening the inhibitor of ALP activity is considered a promising strategy for clinical treatment. Herein, typical inhibitors, such as Na_3_VO_4_ and ATP, were employed to evaluate the screening performance of the *Pf*Ago-based biosensor. The relative activity of ALP gradually decreased with the addition of Na_3_VO_4_ and ATP, and the calculated half maximal inhibitory values were 79.61 and 25.69 μmol L^−1^, respectively, which are consistent with those of previous works(Sun et al. 2018, Zhou et al. 2018). These results reveal that the proposed method could be used to evaluate the inhibition efficiency and screen the ALP inhibitor, suggesting its potential applications in the development of disease therapy tools.

The practical application performance of the *Pf*Ago-based method was verified for ALP activity analysis in human serum (Table 2). The method obtained a similar detection result, with 89%–93.33% recovery, as the clinical assay method on the basis of the ALP-catalyzed colorless *p*-nitrophenyl phosphate substrate into yellow *p*-nitrophenol(Breslow and Katz 1968). Subsequently, various concentrations of ALP were then added to these samples, and the ALP activity of the mixture was analyzed by the *Pf*Ago-based biosensor. Recovery of 94.27% to 103.45% was achieved. These results suggest good accuracy, indicating potential application in clinical diagnosis.

**Table 2.**
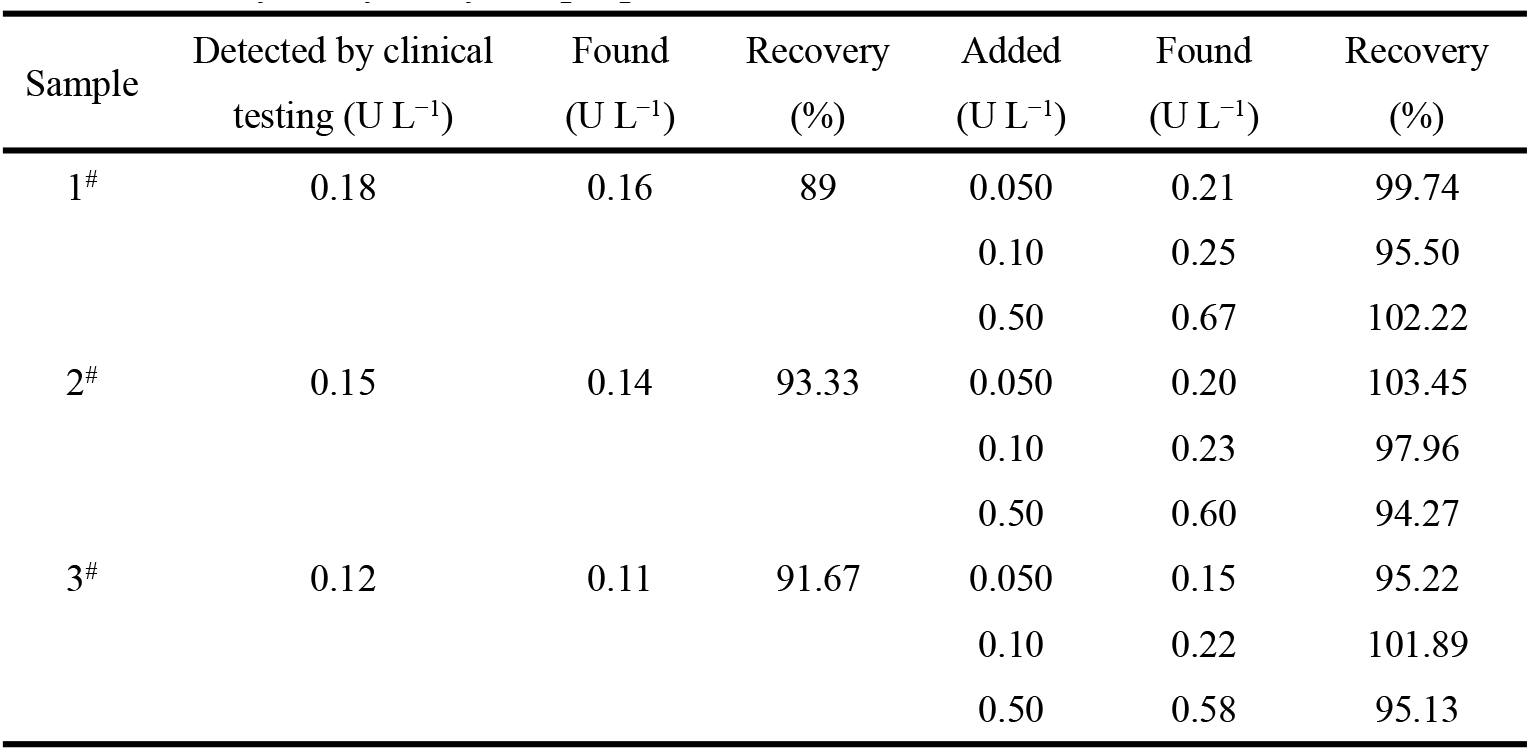
ALP activity analysis by the proposed method.

| Sample | Detected by clinical testing ($\text{U L}^{-1}$) | Found ($\text{U L}^{-1}$) | Recovery (%) | Added ($\text{U L}^{-1}$) | Found ($\text{U L}^{-1}$) | Recovery (%) |
| --- | --- | --- | --- | --- | --- | --- |
| 1<sup>#</sup> | 0.18 | 0.16 | 89 | 0.050 | 0.21 | 99.74 |
|  |  |  |  | 0.10 | 0.25 | 95.50 |
|  |  |  |  | 0.50 | 0.67 | 102.22 |
| 2<sup>#</sup> | 0.15 | 0.14 | 93.33 | 0.050 | 0.20 | 103.45 |
|  |  |  |  | 0.10 | 0.23 | 97.96 |
|  |  |  |  | 0.50 | 0.60 | 94.27 |
| 3<sup>#</sup> | 0.12 | 0.11 | 91.67 | 0.050 | 0.15 | 95.22 |
|  |  |  |  | 0.10 | 0.22 | 101.89 |
|  |  |  |  | 0.50 | 0.58 | 95.13 |

## Conclusion

A *Pf*Ago-based biosensor is proposed for the first time for ALP activity analysis. The proposed biosensor realizes rapid and highly sensitive ALP activity analysis with a LOD of 0.0013 U L^−1^ (3σ/k) and a detection range of 0.0025 U L^−1^ to 1 U L^−1^ within 90 min. The analytical performance of the proposed method is superior to most of the reported methods for ALP activity detection because of the dual signal amplification mode of the *Pf*Ago-mediated multiple-turnover cleavage activity and the pre-amplification technology. The *Pf*Ago-based biosensor has several advantages, including rapidity, high sensitivity, low cost, and sustainability, and potential application in clinic diagnosis and disease therapy.

## Acknowledgments

This work was supported by the National Natural Science Foundation of China (grant 22274060) and a project funded by China Postdoctoral Science Foundation (2022M721269). Part of the work was supported by the Postdoctoral Creative Research Positions of Hubei Province of China (2022). We are grateful for the support of the BaiChuan Fellowship of College of Life Science and Technology, Huazhong Agricultural University.

## Notes

### Competing Interest Statement

The authors have declared no competing interest.

