## Supporting Information for "*Pf*Ago-based dual signal amplification biosensor for rapid and highly sensitive detection of alkaline phosphatase activity"

ORCID:

Yonggang Hu: http://orcid.org/0000-0002-3337-4223.

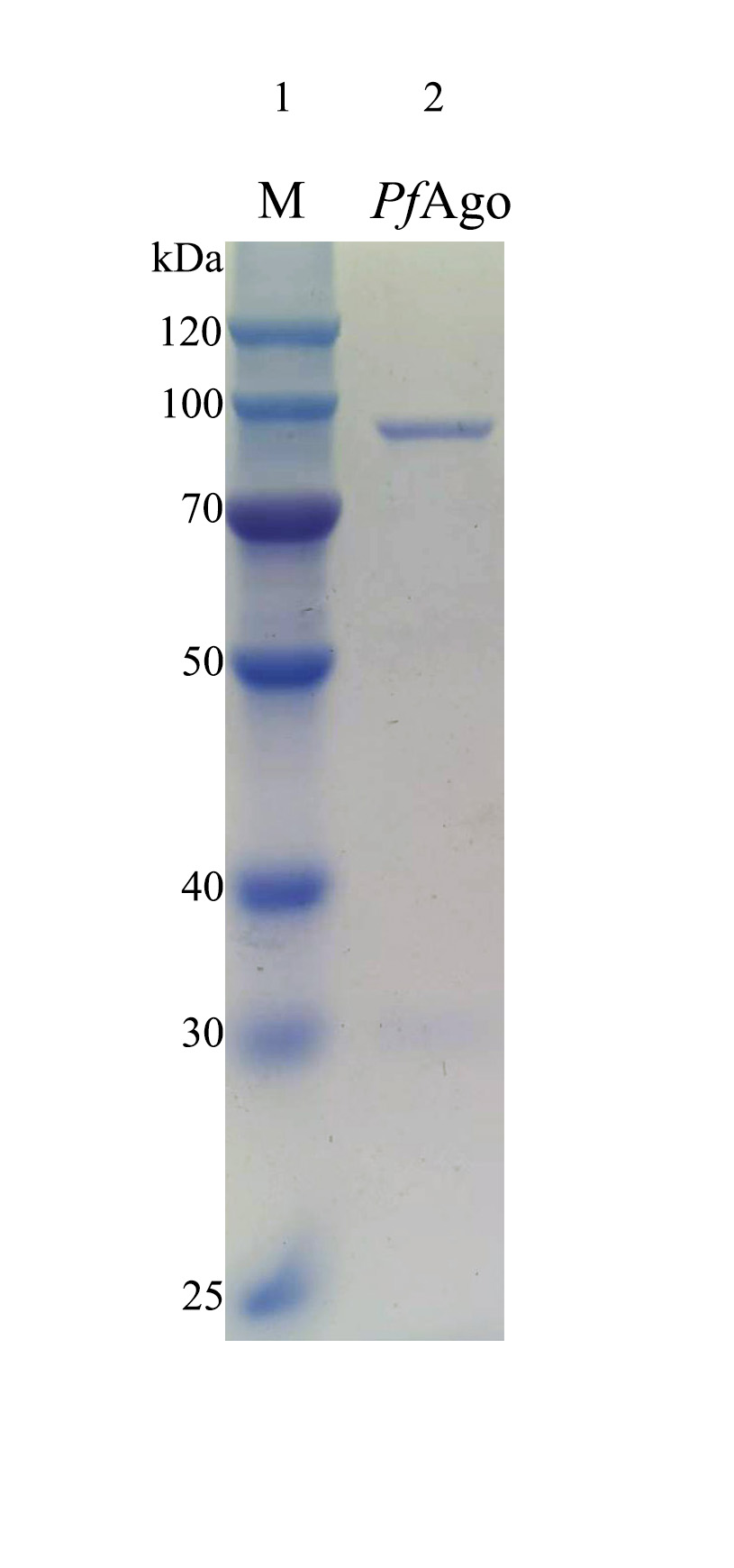

**Figure S1.** SDS-PAGE analysis of the *Pf*Ago protein. (1) Protein marker (M); (2) *Pf*Ago.

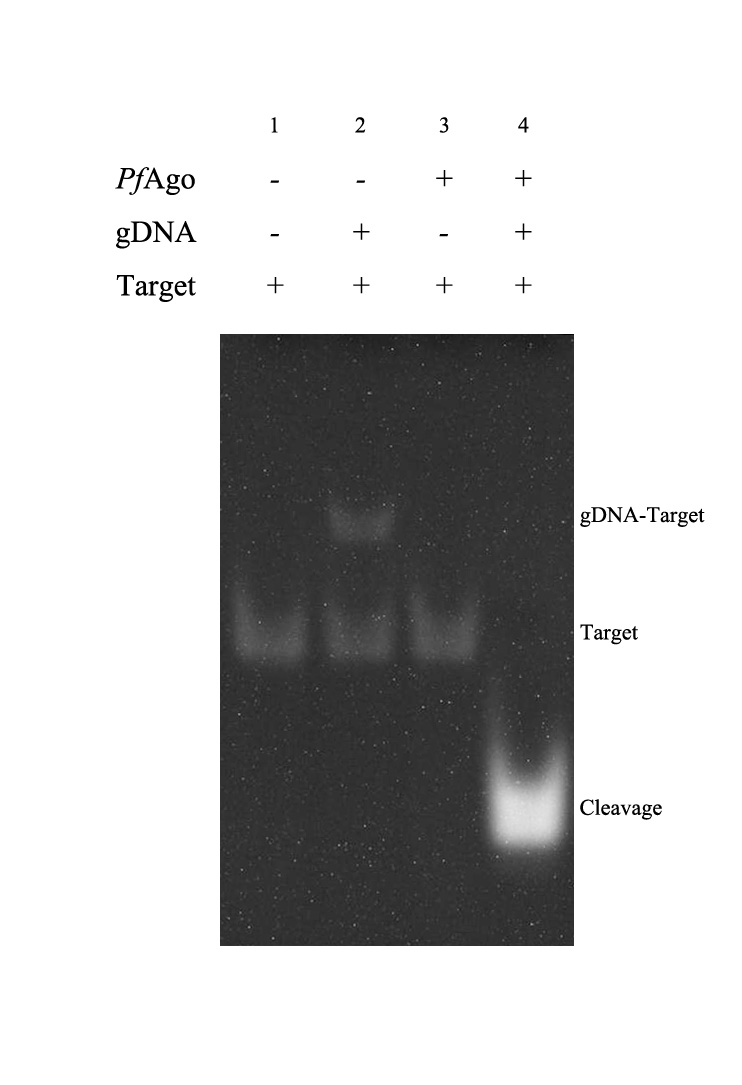

**Figure S2.** PAGE analysis of the *Pf*Ago cleavage activity. (1) ssDNA Target (0.80 μmol L^−1^); (2) gDNA (0.05 μmol L^−1^) + ssDNA Target (0.80 μmol L^−1^); (3) *Pf*Ago (0.08 mg mL^−1^) + ssDNA Target (0.80 μmol L^−1^); (4) *Pf*Ago (0.08 mg mL^−1^) + gDNA (0.05 μmol L^−1^) + ssDNA Target (0.80 μmol L^−1^).

PAGE analysis showed that a band appeared, which corresponds to Target (Lane 1). A higher band appeared which is the dsDNA formed by the complementary base pairing between gDNA and the Target (Line 2). No obvious band change was observed with the addition of the *Pf*Ago without gDNA (Lane 3). An obvious cleavage band appeared with the addition of the gDNA and *Pf*Ago (Lane 4), indicating that *Pf*Ago employs gDNA to form the gDNA-*Pf*Ago complex to activate the cleavage of the complementary ssDNA Target through base pairing.

### Author Contributions

**Weikang Ke，contributor roles:** Data curation: Equal; Formal analysis: Equal; Validation: Equal; Visualization: Equal; Writing – review & editing: Equal

**Yuqing Qin，contributor roles:** Data curation: Equal; Formal analysis: Equal; Funding acquisition: Equal; Methodology: Equal; Supervision: Equal; Visualization: Equal; Writing – original draft: Equal

**Bosheng Zhou，contributor roles:** Conceptualization: Equal; Data curation: Equal; Project administration: Equal; Validation: Equal

**Yonggang Hu，contributor roles:** Funding acquisition: Equal; Methodology: Equal; Project administration: Equal; Resources: Equal; Supervision: Equal
